# Gene panel design for spatial transcriptomics with prioritized gene sets

**DOI:** 10.1101/2022.09.25.509418

**Authors:** Mashrur Ahmed Yafi, Md. Hasibul Husain Hisham, Francisco Grisanti, Atif Rahman, Md. Abul Hassan Samee

**Affiliations:** Department of Computer Science and Engineering, Bangladesh University of Engineering and Technology, Dhaka, 1205, Bangladesh; Department of Integrative Physiology, Baylor College of Medicine, Houston, TX 77030, USA

## Abstract

A fundamental limitation of the emerging single-cell spatial transcriptomics (sc-ST) technologies is their panel size. Being based on fluorescence *in situ* hybridization, an sc-ST dataset can profile only a pre-determined panel of a few hundred genes. This often forces biologists to build panels from only the marker genes of different cell types and forgo other genes of interest, *e*.*g*., genes encoding ligand-receptor complexes or genes in specific pathways. We propose scGIST– a deep neural network that designs sc-ST panels through constrained feature selection. On four datasets, scGIST outperformed alternative methods in terms of cell type detection accuracy. Moreover, unlike other methods, scGIST allows genes of interest to be prioritized for inclusion in the panel while staying within the its size constraint. We demonstrate through diverse use cases that scGIST includes large fractions of prioritized genes without compromising cell type prediction efficacy making it a valuable addition to sc-ST’s algorithmic toolbox.

## Introduction

Spatial transcriptomics (ST) is an emerging suite of technologies that quantify spatially resolved gene expression from intact tissue sections^1–4^. Commercially available ST technologies fall into two types^1–4^. The spot-based ST (spot-ST) technologies quantify gene expression in spatially resolved spots where each spot captures about ten cells. These technologies are sequencing-based and, in principle, capture the transcriptome at each spot. However, the lack of single-cell resolution makes spot-ST data challenging to use in single-cell focused studies, *e*.*g*., to analyze the spatial variation in ligand-receptor usage of a given cell-type^5^.

In contrast to spot-ST, the alternative single-cell ST (sc-ST) technologies capture data from individual cells while recording their locations in the tissue section^3^. However, being FISH-based (Fluorescence In Situ Hybridization), each sc-ST dataset is limited to a pre-determined panel of 10–250 genes^6–12^. This constraint on panel size often limits the potential use of sc-ST data. One primarily prioritizes the marker genes of different cell types in the panel so that the dataset reveals a spatial map of those cell types. However, the panel forgoes other genes of interest relevant to the biological question, *e*.*g*., receptors-ligands or genes in a specific pathway. Thus, there is a critical need for algorithms to design sc-ST gene panels that: (a) accommodate genes of interest beyond cell type markers while (b) accurately detecting different cell types and (c) remaining within the panel’s size limit. Importantly, these algorithms should be designed to deal with constraints that biologists face in their practical applications. For example, when a biologist wishes to set differential priorities to the genes of interest, the algorithm should maximize the inclusion of higher priority genes. Similarly, when a biologist wants to include protein complexes (such as ligand-receptor complexes) in the panel, the algorithm should aim at including all genes encoding the proteins in a complex. To address these versatile needs while ensuring cell type detection accuracy, here we present scGIST (single cell Gene-panel Inference for Spatial Transcriptomics).

scGIST is a deep neural network with a custom loss function that casts sc-ST panel design as a constrained feature selection problem^13^. Given a single-cell (sc) RNA-seq dataset annotated for cell types, scGIST learns to classify the individual cells given their gene expression values. Notably, its custom loss function aims at maximizing both cell type classification accuracy and the number of genes included from a given gene set of interest while staying within the panel’s size constraint. Thanks to this loss function, scGIST can cater to a wide range of applications and enable gene panel design to address specific research questions. For example, as an input to the model, a biologist can specify differential priorities for genes in the given gene set. Such scenarios may arise when they wish to assign higher priorities to genes for which FISH probe design is easy or to genes that are expressed above a threshold level. In such cases, scGIST will try to maximize the number of highest priority genes in the panel without compromising cell type prediction accuracy to a great extent. scGIST also allows users to specify a list of genes of interest or protein complexes, such as ligand-receptor complexes. For every complex, the model will aim at selecting the complete set of genes encoding the proteins in that complex.

We demonstrate scGIST’s efficacy and versatility by bench-marking it on four datasets comprising 2700–20000 cells and 8–23 cell types, and through a number of use cases. We are unaware of other methods with such generic applicability as scGIST. Three recent sc-ST marker selection methods, namely scGeneFit^14^, geneBasis^15^, and SMaSH^16^, can be used to identify marker genes for a given panel size. scGeneFit^14^ selects marker genes given scRNA-seq data and cell labels using linear optimization techniques. geneBasis^15^ designs panels by greedily adding a new gene which captures the maximum Minkowski distance between true manifold and current manifold. It allows a priori addition of genes of interest to the panel but, unlike scGIST, cannot trade-off their inclusion with accuracy. SMaSH^16^ is a machine learning framework which learns a classifier for cell types and then greedily selects markers using various criteria.

We first compare scGIST with these three methods^14–16^ in a setting with no prioritized genes. Our benchmarking demonstrates that scGIST consistently outperforms the alternative methods and the margins are remarkable for small panel sizes (*<* 50). Next we show that scGIST can include other genes or complexes of interest such as genes with easy to design probes, genes in specific pathways and receptor-ligand complexes in the panel without sacrificing cell type prediction accuracy substantially. Overall, with its high accuracy and versatile applicability, scGIST addresses a critical need in current sc-ST methods.

## Results

### A deep learning model for marker gene selection

We solve the gene panel selection problem by formulating it as a feature selection problem for multi-class classification in neural networks^13^. Our method, illustrated in Figure 1A, takes as input gene expression values from an scRNA-seq experiment, labels for the cells, and a target gene panel size. The cell labels may represent cell states or cell types, such as the tissue or the organ they originated from. If the labels are not available, they may be obtained by clustering, or through other approaches^17^. It then generates as output a gene panel that can accurately distinguish among cell types. These genes can subsequently be used as marker genes for technologies such as spatial transcriptomics.

**Figure 1.**
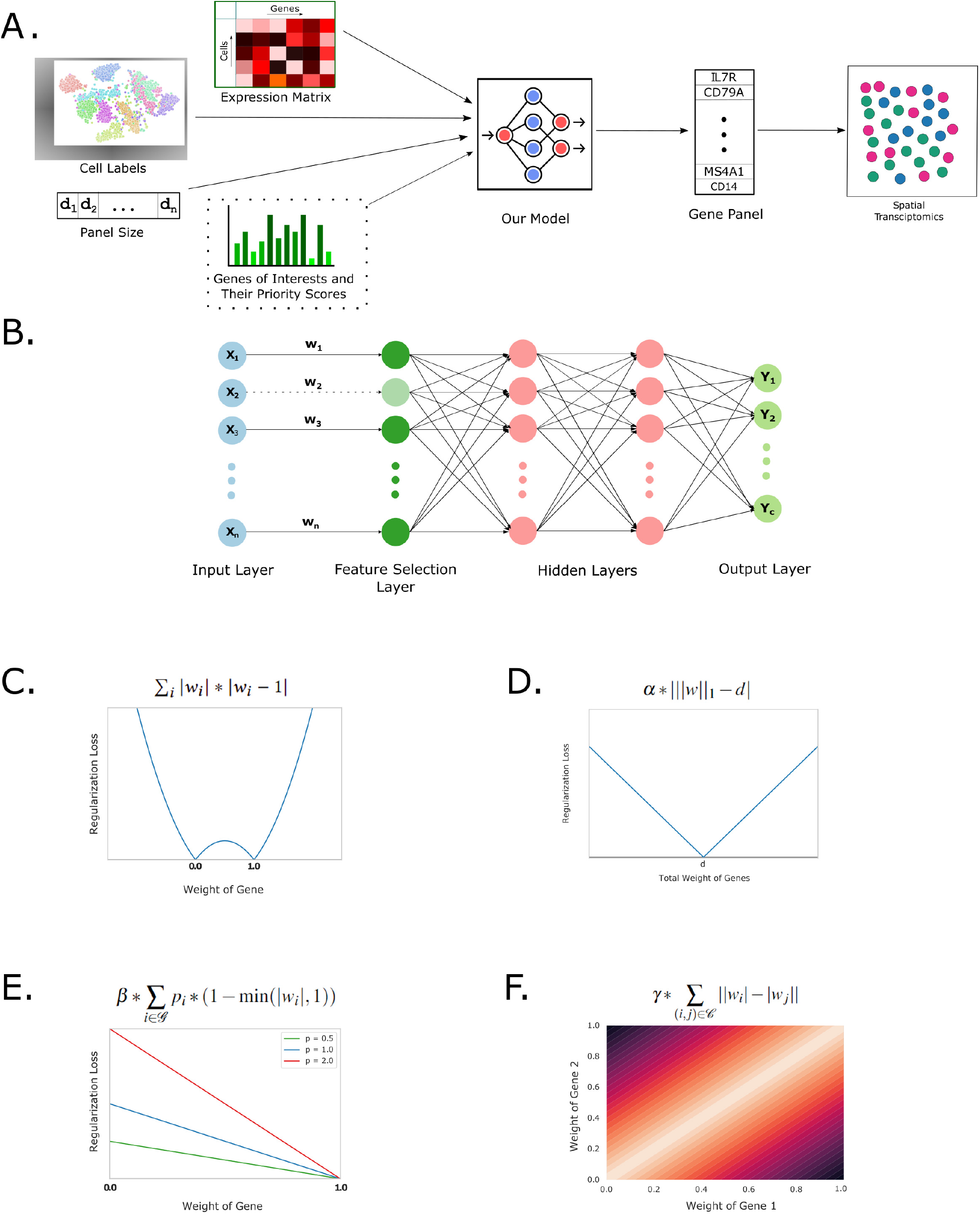
Overview of scGIST. **A**. scGIST takes as input expression values from an scRNA-seq experiment, labels for the cells, a target gene panel size, and optionally a list of genes of interest and their priority scores. It uses a deep learning model to learn a set of genes to predict cell types which can be used as a gene panel for single cell spatial transcriptomics assays. **B**. The neural network model of scGIST consists of a one-to-one feature selection layer connected to the input layer. This is then followed by two fully connected layers and finally an output layer. The model has a custom loss function **C**. to drive weights in the feature selection layer towards 0 or 1, **D**. to select the number of genes equal to the panel size (*d*), **E**. to prioritize genes of interest according to user provided priority scores (*p*_*i*_), and **F**. to select both or none of the genes from specified complexes.

At the core of our method, there is a deep learning model for multi-class classification with a custom loss function for feature selection. The inputs to this neural network are the log-normalized counts of the scRNA-seq data and the outputs are the probabilities of each type of cells. The neural network, shown in Figure 1B, contains a one-to-one linear layer (feature selection layer) between the input layer and the first fully connected layer, which is then followed by 2 fully connected layers. Finally, there is an output layer which predicts the labels of the cells. To perform feature selection, we use a custom loss function^13^ which includes the following components (see Methods for details):

- A regularization term in the one-to-one linear layer that drives the weights of each gene towards either 0 or 1 (Figure 1C). This term promotes sparsity of the weights. Additionally, a regularization term corresponding to all other layers ensures that no small weight in the one-to-one layer is amplified by any of the subsequent layers.
- A penalty term that comes into effect if the sum of the weights of the one-to-one layer is not equal to the target panel size *d* (Figure 1D). Since these weights are close to zero or one, this sum represents the number of features taken. This term can also optionally be set to penalize only if the sum of the weights exceeds *d* (Supplementary Note 3.1).

The model is then trained using the provided scRNA-seq expression values and the cell labels, and the genes corresponding to the *d* highest weights in the first layer are selected as the gene panel.

### Regularization for inclusion of other genes and complexes of interest

In addition to marker gene selection, scGIST allows users to provide as option a list of genes of interest and their priority scores. These genes may represent known marker genes, genes with easy to design probes as well as other genes and complexes of interest. The method places a greater emphasis on inclusion of the user specified genes based on their priority scores. This is done as follows (details in Methods):

- scGIST may be provided with lists of genes of interest to be included in the panel along their priority scores optionally. For each user specified gene, a penalty proportional to its priority score is added if it is not selected in the panel (Figure 1E).
- Users may also list receptor-ligand or other complexes for inclusion in the gene panel. For each pair of genes in the complexes, a penalty term proportional to the difference of their weights in the one-to-one layer is added to the loss function (Figure 1F). As a result, inclusion of only a subset of the genes in a complex will result in an additional cost.

To assess the performance of scGIST, we use it to analyze four real datasets - peripheral blood mononuclear cells (PBMC 3k) (https://support.10xgenomics.com/single-cell-gene-expression/datasets/1.1.0/pbmc3k), Head and Neck Cancer^18^, Tabula Sapiens^19^, and Mouse Endoderm^20^. First, we present a comparison of its performance with those of existing gene panel selection tools scGeneFit, geneBasis and SMaSH, and subsequently analyze its effectiveness in including user provided genes and complexes of interest.

### Gene panel selection for cell type identification

We ran scGIST as well as scGeneFit, geneBasis and SMaSH on the four aforementioned datasets for varying panel sizes and calculated their accuracy (Figure 2A and Supplementary Tables 1,3,5,7). The values for scGIST are averages over five runs. We observe that our method performs as well as the state-of-the-art methods in the PBMC 3K dataset and outperforms all methods in the other three datasets. scGIST often outperforms other methods for marker selection substantially, which is especially noticable for small panel sizes. As an example, for the Head and Neck Cancer dataset, scGIST has an accuracy of 95.31% at panel size of 50 whereas scGeneFit, geneBasis and SMaSH have accuracy of 77.18%, 82.67% and 84.25% respectively. For smaller panel sizes, the differences are even more pronounced.

**Figure 2.**
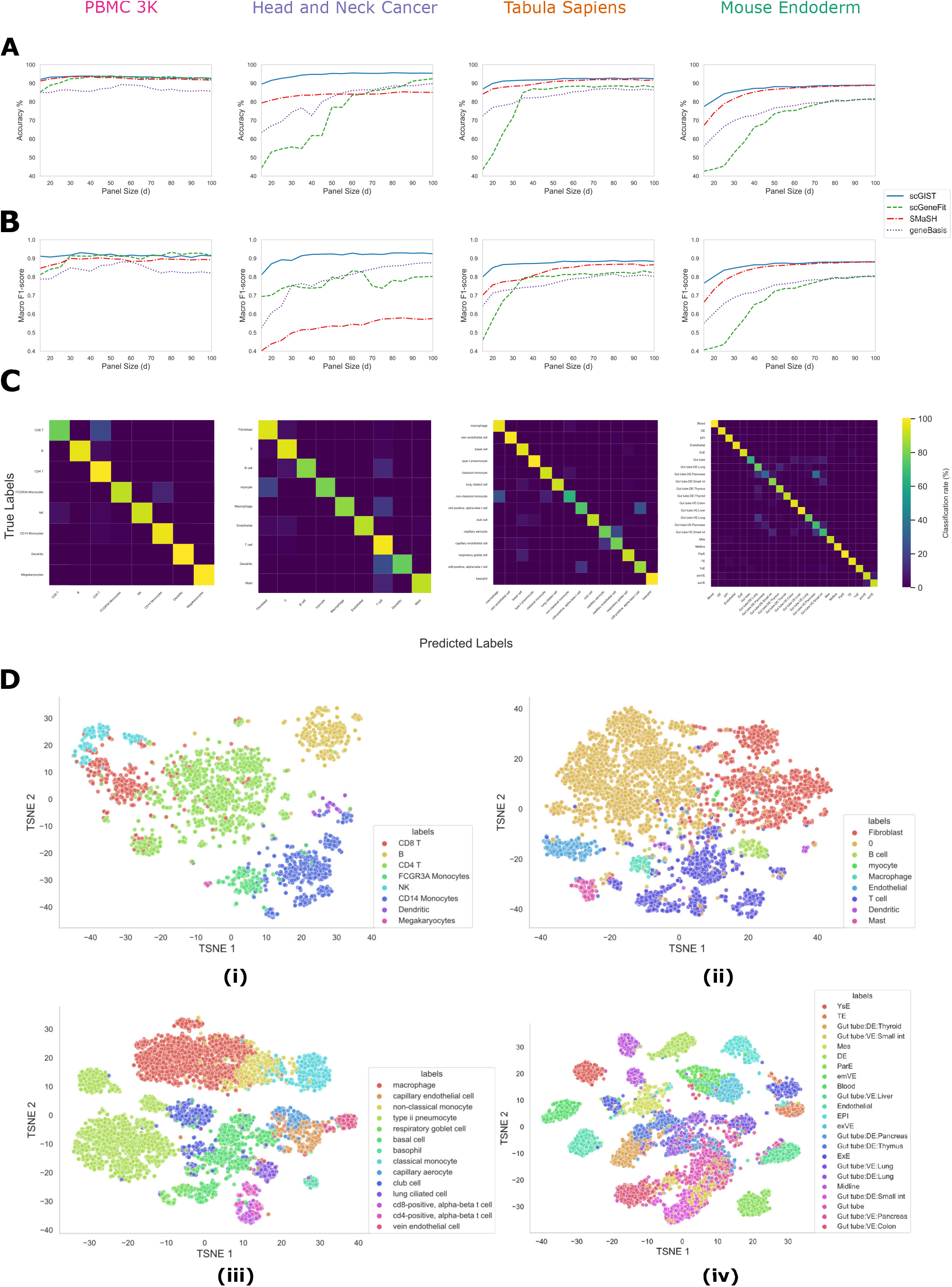
Performance of various methods on four different datasets. **A**. Accuracy for variable panel sizes, **B**. Macro F1-scores for variable panel sizes, **C**. Confusion matrices for scGIST for panel size = 60, **D**. t-SNE plots using the gene panel of size 60 selected by scGIST for (i) PBMC 3K, (ii) Head and Neck Cancer, (iii) Tabula Sapiens, and (iv) Mouse Endoderm datasets.

Macro F1 scores (Supplementary Note 3.2) were also computed for the methods for different panel sizes as accuracy can be misleading when there is class imbalance. We find that scGIST outperforms other methods in terms of macro F1 score as well (Figure 2B and Supplementary Tables 2,4,6,8). It is worth noting that although SMaSH has more than 80% accuracy in the Head and Neck Cancer dataset, it has low macro F1-score in that dataset. For example, it has macro F1-score of 0.58 despite having an accuracy of 85.06% at panel size of 100. The high macro F1 scores of our method is due to its accuracy across all cell types which is illustrated in the confusion matrices in Figure 2C.

We also created t-SNE plots^21^ for visualization of the classification into various cell types (Figure 2D). For each of the four datasets, 60 marker genes were first selected by our method and then t-SNE plots were created using those genes. It can be observed from the plots that the genes selected by our method can largely separate the different cell types in all datasets. It is worth noting that all four methods perform poorly in distinguishing various gut tube cell types in the Mouse Endoderm dataset compared to all other cell types in all datasets.

### Performance on rare cell types

As indicated by the macro F1-scores, scGIST performs well on rare cell types. Figure 2C and Supplementary Figures 2-5 show the cell type confusion matrices of scGIST as well as scGenefit, geneBasis and SMaSH for the four datasets for panel size of 60. Cell type abundances are shown in ordered bar plots in Supplementary Figure 1. We observe that scGIST, scGeneFit and geneBasis show consistent performance across all cell types with scGIST outperforming the other two overall. However, SMaSH often demonstrate poor performance for rare cell types. As an example, SMaSH is unable to correctly predict any of the dendritic cells, the second rarest cell type in the PBMC 3k dataset whereas scGIST, scGeneFit and geneBasis correctly predicted 100%, 89% and 78% of them respectively.

### Prioritization of genes with easy to design probes

scGIST provides the option to prioritize various genes and complexes of interest, such as genes for which probes are easy to design, through user given priority scores. To explore this, we experiment with the Mouse Endoderm dataset. Lohoff *et al*.^22^ reported 387 target genes for which probes are easier to design and are good at cell type classification for the single-cell molecular map of mouse gastrulation and early organogenesis^23^. Out of these 387 genes, 2 were dropped at the preprocessing step. The remaining 385 genes were fed to our model with priority scores set to 0 and 1 in two experiments.

Figure 3A and Figure 3B show the accuracy of the method and the numbers of easy to probe genes included in the panel for priority scores of 0 and 1 respectively for various panel sizes. We observe that when the priority scores are set, many more easy to probe genes are included in the panel without much reduction in accuracy. The actual numbers are listed in Supplementary Tables 9-11. For example - for panel size 100, when no priority score is set, 89.26% accuracy is achieved while including 36 genes with easy to design probes. But when the priority scores are set to 1, 60 such genes are included with accuracy only dropping slightly to 88.64%. Even if we consider a large panel of size of 500, we find that only 133 genes with easy to design to probes are included in the panel when no priority is set whereas all 385 are selected when priority scores are set to 1.

**Figure 3.**
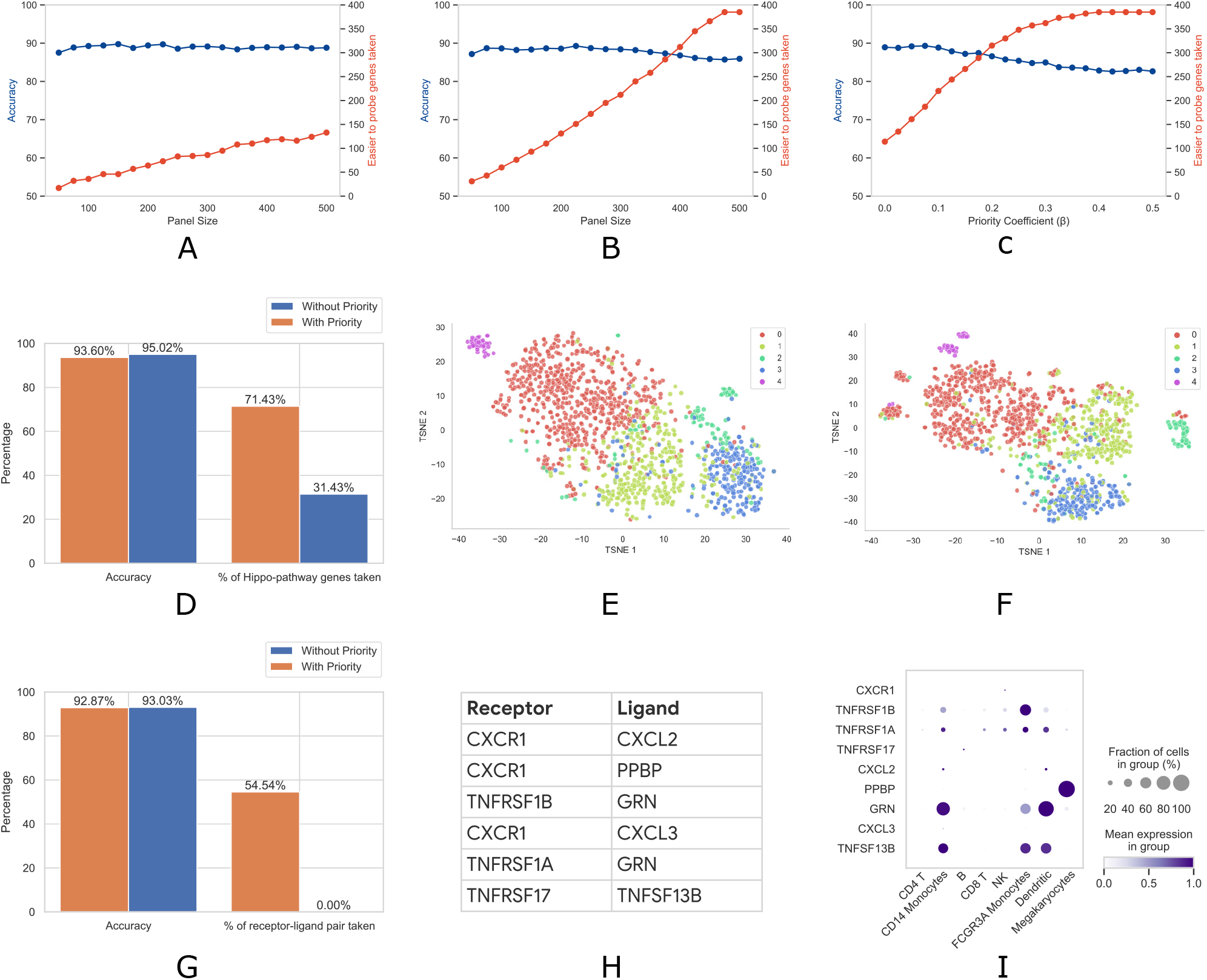
**A-C**. Analysis of prioritization of easy to probe genes for the Mouse Endoderm dataset. Accuracy and the numbers of easy to probe genes included in gene panels when **A**. priority scores of all genes = 0, and **B**. priority scores of easy to probe genes = 1, co-efficient of priority (*β*) = 0.2 for varying panel sizes, and **C**. priority scores of easy to probe genes = 1, panel size = 500 for varying co-efficient of priority (*β*). **D-F**. Hippo-pathway gene analysis on the Head and Neck Cancer dataset. **D**. Comparison of accuracy and number of Hippo pathway genes taken in a panel of size 60 with and without prioritization. t-SNE plots of Fibroblast subtypes using gene panel of size 60 with **E**. Hippo-pathway genes prioritized and **F**. Hippo-pathway genes not prioritized. **G-I**. Receptor-ligand complex analysis on the PBMC 3k dataset. **G**. Comparison of accuracy and number of receptor-ligand pairs taken in a panel of size 60 with and without prioritization. **H**. Receptor-ligand complexes included in the panel. **I**. Dot plot showing, for each gene encoding receptors or ligands, fractions of cells of each type where they are expressed, and their mean expressions.

The importance of the prioritized genes can also be tuned through the co-efficient of priority (*β*). Figure 3C and Supplementary Table 11 show the accuracy and the numbers of easy to probe genes included in the panels against varying co-efficient of priority (*β*) for panel size of 400. For *β* = 0, which is equivalent to not prioritizing any gene, we find that 114 of those genes are included in the panel with cell type prediction accuracy of 88.93%. For the default *β* = 0.2, the number of such genes in the panel increases to 315 with accuracy dropping to 86.57%. Finally, for *β ≥*0.4, all 385 easy to probe genes are selected in the gene panel.

### Inclusion of other genes interest (Hippo pathway genes)

In many experiments, genes of interest, other than the marker genes to identify cell types, need to be included in the panel to address specific biological questions. One approach to this may be to include such genes in the panel first and then to select marker genes to fill the remaining slots in the panel. However, this approach may lead to drastic reduction in cell type prediction accuracy. scGIST provides the option to systematically trade-off between genes-of-interest inclusion and prediction accuracy through incorporation of user provided priority scores in the loss function.

As a case study, we consider the role of genes in the Hippo signaling pathway in fibroblast cells in the Head and Neck Cancer dataset. Hippo pathway genes have been conjectured to play roles in fibroblast cell differentiation^24^, cancer development^25,26^, etc. and as such, it may be of interest to include them in the panel. To analyze this, we assign all 35 genes known to be involved in the Hippo signalling pathway priority scores of 0 and 1 in two experiments. All other genes were given priority scores of 0 in both experiments. We then design panels of size 60 for the Head and Neck Cancer dataset. The accuracy and the percentages of Hippo pathway genes included in the panel in the two settings are shown in Figure 3D. We find that when no priority is provided, 31.43% of Hippo pathway genes are included in the panel and the prediction accuracy is 95.02% for all cell types in the Head and Neck Cancer dataset. However, when they are assigned priorities of 1, 71.43% of those genes are selected and the accuracy only slightly decreases to 93.60%.

Next we isolate the fibroblast cells in the dataset and identify subtypes within them. We perform principal component analysis (PCA) on the expression values of all genes and construct a graph based on the first 60 principal components. We then run the community detection algorithm Leiden^27^ on this graph to cluster the cells which reveals four subtypes (Types 0-3). Subsequently, we create a t-SNE plot using the 60 genes selected in the panel when the Hippo pathway genes are prioritized which is shown in Figure 3E. We observe that a subset of cells appear to form a separate cluster (labelled Type 4 and shown in a different color). This subset of cells consistently shows up as a different cluster in t-SNE plots with various settings (not shown). However, when a similar t-SNE plot is constructed using the gene panel with Hippo pathway genes not prioritized, this cluster is split into multiple sub-clusters with some cells grouped with Type 0 cells (Figure 3F). This illustrates that inclusion of specific genes of interest may provide insights that may not be obtained from marker genes in the original dataset because expressions of marker genes may not correlate well with that of the genes of interest in question.

### Inclusion of complexes of interest (receptor-ligand complexes)

Finally, scGIST also permits researchers to prioritize genes encoding various protein complexes such receptor-ligand complexes. Cell-cell interactions drive cell differentiation and organ development, and disruption in signalling pathways have been implicated in various diseases^28^. Receptor-ligand complexes are key to understanding intercellular communication and as a result it is important to measure their expression, especially at the surfaces of tissues. However, these genes are often not included in gene panels as they are not markers for cell types or are expressed in a small number of cells.

To analyze this, we collected 11 receptor-ligand complexes from CellPhoneDB^29^. We then encoded them using pairs (see Methods for details) and experimented with the PBMC 3k dataset. The results are summarized in Figure 3G. We observe that the panel includes none of the receptor-ligand pairs when the complexes are not prioritized. However, when they are prioritized, as many as 54.54% of the pairs are selected in the panel. Moreover, this is at the expense of only a small decrease in accuracy from 93.03% to 92.87%.

The receptor-ligand complexes that are included in the gene panel are provided in Figure 3H while a dot plot of expressions of the genes by cell types is shown in Figure 3I. The dot plot illustrates, for each cell type, the fraction of cells where each of those genes are expressed and their mean expressions. We observe that the gene *PPBP* appears to be only expressed in Megakaryocytes and hence a good candidate as a marker. However, some genes such as *TNFRSF1A* and *TNFRSF1B* are expressed in a number of cell types while *CXCR1* is expressed highly in only a small fraction of NK cells. Similarly, *CXCL2* is expressed highly in a small number of CD14 Monocytes and Dendritic cells. scGIST is able to include these genes in the panel based on user prioritization and thus potentially providing insights which would be missed otherwise.

## Discussion

By adopting a powerful neural network architecture for feature selection and a custom loss function, we have shown that scGIST is an effective and broadly applicable tool for sc-ST panel design. The model optimizes cell type detection accuracy while satisfying users’ need regarding a gene set of interest and staying within the size limit of the sc-ST gene panel.

We have demonstrated scGIST’s success on four datasets and under different use cases. While it is difficult to make a theoretical argument explaining this success, we believe that scGIST owes its success as much to its architecture and the loss function as to the inherent covariance structures of gene expression relationships in single-cell data. Recent works from Aviv Regev and colleagues^6,30^ have shown that genes in these datasets form co-expression modules – the expression of any gene from a module is generally sufficient to recover the expression of other genes in that module. We believe that scGIST can successfully leverage this property to select genes satisfying various constraints without sacrificing classification accuracy.

Importantly, scGIST has consistently outperformed the alternative tools^14–16^ that can be used to derive marker gene panels under size constraints. Again, it is difficult to give theoretical arguments that place one method clearly as the best method, but scGIST presumably outperformed scGeneFit^14^ by exploring a larger function space. Although SMaSH^16^ is a deep neural network like scGIST, SMaSH requires posthoc model interpretation methods to rank the features and applies a greedy feature selection strategy. In contrast, scGIST integrates feature ranking and all other constraints into the loss function. We believe, this helps scGIST make the necessary feature trade-offs directly during model optimization and select the most appropriate features that maintain accuracy while satisfying the other constraints as much as possible. In comparison to geneBasis^15^, scGIST does not require greedy feature selection and need not make assumptions about cell-to-cell distance metrics – both characteristics should have helped scGIST to outperform geneBasis.

scGIST has opened up several exciting directions for future research. First, currently all methods, including scGIST, design panels for a single biological context, *e*.*g*., a particular tissue type or a particular timepoint. An impactful direction would be to design panels that maximize the use of the selected genes (*i*.*e*., their probes) for multiple biological contexts. Secondly, now that scGIST enables biologists to specify gene priorities, we anticipate there will be more research on modeling the ease of probe design for given transcript sequences^3^. Finally, our future work will focus on implementing scGIST as an interactive tool for panel design, where the model will explain why it could not satisfy all constraints and what potential trade offs might be necessary to optimize a particular goal, *e*.*g*., including all ligand-receptor complexes.

## Methods

### scGIST model

scGIST takes as input *N* instances {(**x**_**1**_, *y*_1_), (**x**_**2**_, *y*_2_), …, (**x**_**N**_, *y*_*N*_)} where **x**_**i**_ ∈ ℝ^*n*^ is log normalized gene expression values of *n* genes of the *i*-th cell and *y*_*i*_ ∈ {1, 2, …, *c*} is the corresponding cell label. It learns a deep neural network model from the training instances to predict cell labels *y* from gene expression values **x** and uses a special regularization function to ensure that only a subset of the genes are used for prediction^13^.

The deep neural network model is shown in Figure 1B. Let **x** = (*x*_1_, *x*_2_, …, *x*_*n*_) be the inputs to the network. To perform feature selection, we add a one-to-one linear layer following the input layer. If the weights of one-to-one linear layer are **w** = (*w*_1_, *w*_2_, …, *w*_*n*_), then the output of the layer will be **w.x** = (*w*_1_*x*_1_, *w*_2_*x*_2_, …, *w*_*n*_*x*_*n*_). Here **w** has to be a sparse binary vector in order to perform feature selection.

Following the one-to-one layer, there are 2 fully connected layers with 32 and 16 neurons respectively with ReLU activation and L2 regularization functions. Finally, there is an output layer with soft-max activation function having *c* neurons. We denote the parameters of these layers by **W**^(**k**)^ and **b**^(**k**)^ for *k* = 1, 2, 3, where **W**^(**k**)^ is the weight matrix of the layer and **b**^(**k**)^ is the corresponding bias vector. So, the parameters of the model can be denoted together as *θ* = {**w, W**^(**1**)^, **b**^(**1**)^, **W**^(**2**)^, **b**^(**2**)^, **W**^(**3**)^, **b**^(**3**)^ }.

To learn the model parameters, we have to minimize the following loss function:

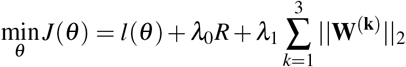

Here, *l*(*θ*) is the log-likelihood of the given data which can be written as

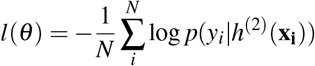

where *h*^(2)^(**x**_**i**_) is the output of the second fully connected layer given input **x**_**i**_. The weights are learnt by minimizing the loss function and the genes corresponding to the *d* highest weights in the feature selection layer are selected as the panel. *R* is a special regularization term in the one-to-one layer that is used for feature selection which is described in the next section. 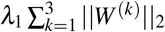 is used for L2 regularization of the final three layers. This is used to reduce bias and to make sure that no small value of *w* in the one-to-one layer is amplified by any of **W**^(**k**)^ in the subsequent layers.

### Regularization

In order to make **w** a sparse binary vector to perform feature selection, and to prioritize inclusion of user specified genes and complexes, we include the following special regularization in the loss function:

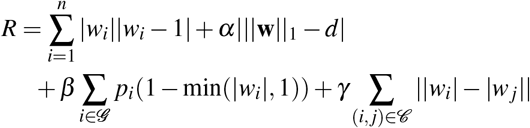

where

1. The first term 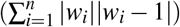 drives the weights *w*_*i*_ towards either 0 or 1. We can see from Figure 1C that the error is minimum (0) when either *w*_*i*_ = 0 or *w*_*i*_ = 1 for *i* = 1, …, *n*. Thus, this regularization term promotes the sparsity of the weights.
2. *α*|||*w*||_1_ −*d*| makes sure that the number of genes used to predict cell labels is equal to *d*, the target gene panel size. As each |*w*_*i*_| is close to 0 or 1, the sum indicates the number of features taken. We will assign a penalty if the number of features taken is more or less than *d*. Here *α* is a hyperparameter that determines how much the model is penalized if it takes more or less than *d* parameters. We have found empirically that setting *α* = 1 provides satisfactory performance. Alternatively, this term can be set in a way to penalize the model only if the number of features exceeds *d* (Supplementary Note 3.1).
3. *β* ∑_*i*∈ 𝒢_ *p*_*i*_(1− min(|*w*_*i*_|, 1)) is an additional term which makes the model prioritize inclusion of genes of interest. Here 𝒢 is the set of user provided genes of interest and *p*_*i*_ is the priority score of the *i*-the gene. If *p*_*i*_ *>* 0, we apply an additional penalty if that gene is not included in the panel (Figure 1E). The amount of penalty is proportional to the priority of the gene. Here *β* indicates how much significance we put on selecting the genes of interest. There is a trade-off between accuracy and how many genes of interest are taken because these genes may not provide much information for classification. We found that setting *β* = 0.5 provides a good balance.
4. Finally, the term *γ* ∑_(*i,j*)∈𝒞_ ||*w*_*i*_| − |*w*_*j*_|| is used for inclusion of complexes. Here 𝒞 = {(*i*_1_, *j*_1_), (*i*_2_, *j*_2_), …} is a set of pairs of genes that represents complexes of interest. These complexes can be provided by users to indicate that either both genes from a pair or none should be selected. The regularization function will assign a penalty if two genes of a pair do not have the same weights. By default, one of the two genes in the pair is randomly assigned a priority score of 1. However, the user has the option of assigning priority scores to both of the genes. If there are complexes with more than two interacting genes, they can be encoded by enumerating all pairs in the complex.

### Datasets

To evaluate our method, we have used the following four datasets:

### PBMC3k

This is a dataset of peripheral blood mononuclear cells (PBMCs) from a healthy donor freely available from 10x genomics (https://support.10xgenomics.com/single-cell-gene-expression/datasets/1.1.0/pbmc3k). PBMCs are primary cells with relatively small amounts of RNA (∼1pg RNA/cell). There are 2,700 single cells that were sequenced on the Illumina NextSeq 500. The number of genes is around 1800 after preprocessing. There are 8 cell types present in the dataset.

### Head and Neck Cancer

The dataset contains transcriptomes of 6000 single cells from 18 head and neck squamous cell carcinoma (HNSCC) patients, including five matched pairs of primary tumors and lymph node metastases which is collected by Puram SV, et al.^18^ There are 9 cell types present with imbalanced distribution.

### Tabula Sapiens (Lungs)

The Tabula Sapiens Dataset^19^ is a human reference atlas which contains around 500,000 cells from 24 different tissues and organs, many from the same donor. This atlas enabled molecular characterization of more than 400 cell types, their distribution across tissues and tissue specific variation in gene expression. We have used a subset of this dataset consisting of cells only from lungs which contains 14 cell types.

### Mouse Endoderm Datase

This is a dataset of single-cell transcriptomes representing all endoderm populations within the mouse embryo until midgestation to delineate the ontogeny of the mammalian endoderm^20^. The dataset contains more than 110,000 single-cell transcriptomes. We have used a subset of this dataset containing 20,000 cells and 23 cell types.

### Data preprocessing

All the datasets were preprocessed using SCANPY^31^ by following a standard pre-processing workflow for scRNA-seq data in Seurat^32^. First, a basic filtering was applied where any cell with less than 200 genes or any gene expressed in less than 3 cells were discarded. Total-count normalization and logarithmization was then performed on the datasets. Highly variable genes were subsequently identified and rest of the genes were discarded. Effects of total counts per cell and the percentage of mitochondrial genes expressed were regressed out. Each gene expression was finally scaled to unit variance and values exceeding standard deviation 10 were clipped at In addition, a maximum of 1000 cells per cell type were retained.

### Evaluation

To evaluate the performance of different marker gene panel selection methods, we followed the approach used to evaluate SMaSH^16^. each of them was executed to find a marker gene panel of size *d*. Only the selected genes were kept in the dataset i.e. all other genes that are not present in the marker gene panel were removed from the dataset. So the new dataset was of dimension *n*× *d* where *n* is the number of cells in the dataset. The new dataset was split into train and test set in 75:25 ratio. Then a k-nearest neighbor (k-NN) classifier was trained using the training dataset. After that accuracy and macro F1 score (Supplementary Note 3.2) was measured on the test dataset. This accuracy and macro F1-score are used in this paper to compare different methods.

## Code Availability

scGIST is implemented in Python and is freely available for download. The code as well as scripts to reproduce the results are available at https://github.com/yafi38/scGIST.

## Data Availability

The datasets used in this paper are publicly available. The download information is as follows:

- PBMC 3k: Available for download through the 10x Genomics website.
- Head and Neck Cancer^18^: Available through the Gene Expression Omnibus, under accession number GSE103322.
- Tabula Sapiens^19^: Available for download through the Tabula Sapiens website.
- Mouse Endoderm^20^: Available through the Gene Expression Omnibus, under accession number GSE123046.

## Supplementary information

### 1 Supplementary Figures

**Figure S1:**
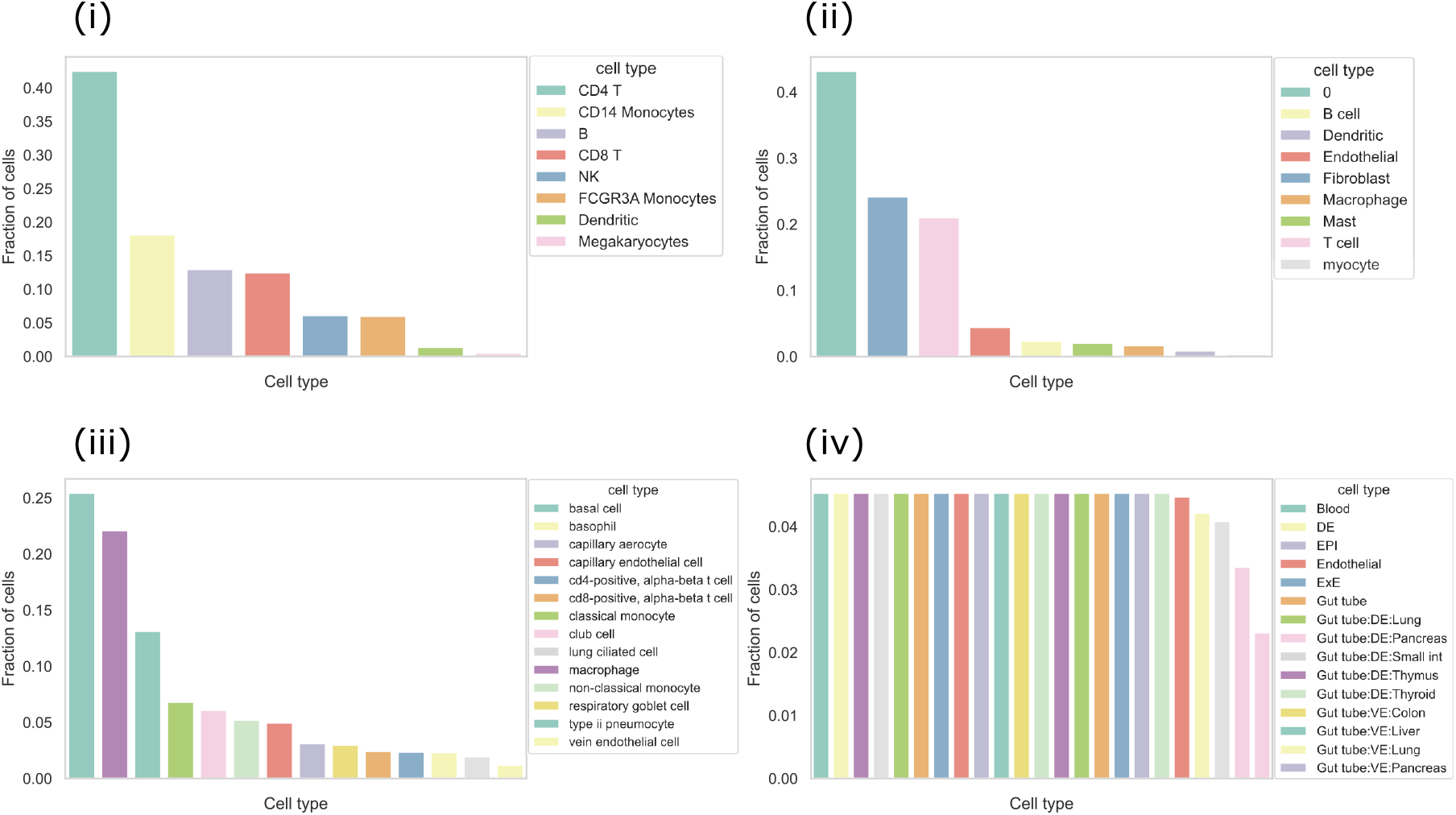
Ordered bar plot representing relative cell type abundance in different dataset. Panel **(i)** PBMC 3K, **(ii)** Head and Neck Cancer, **(iii)** Tabula Sapiens, **(iv)** Mouse Endoderm.

**Figure S2:**
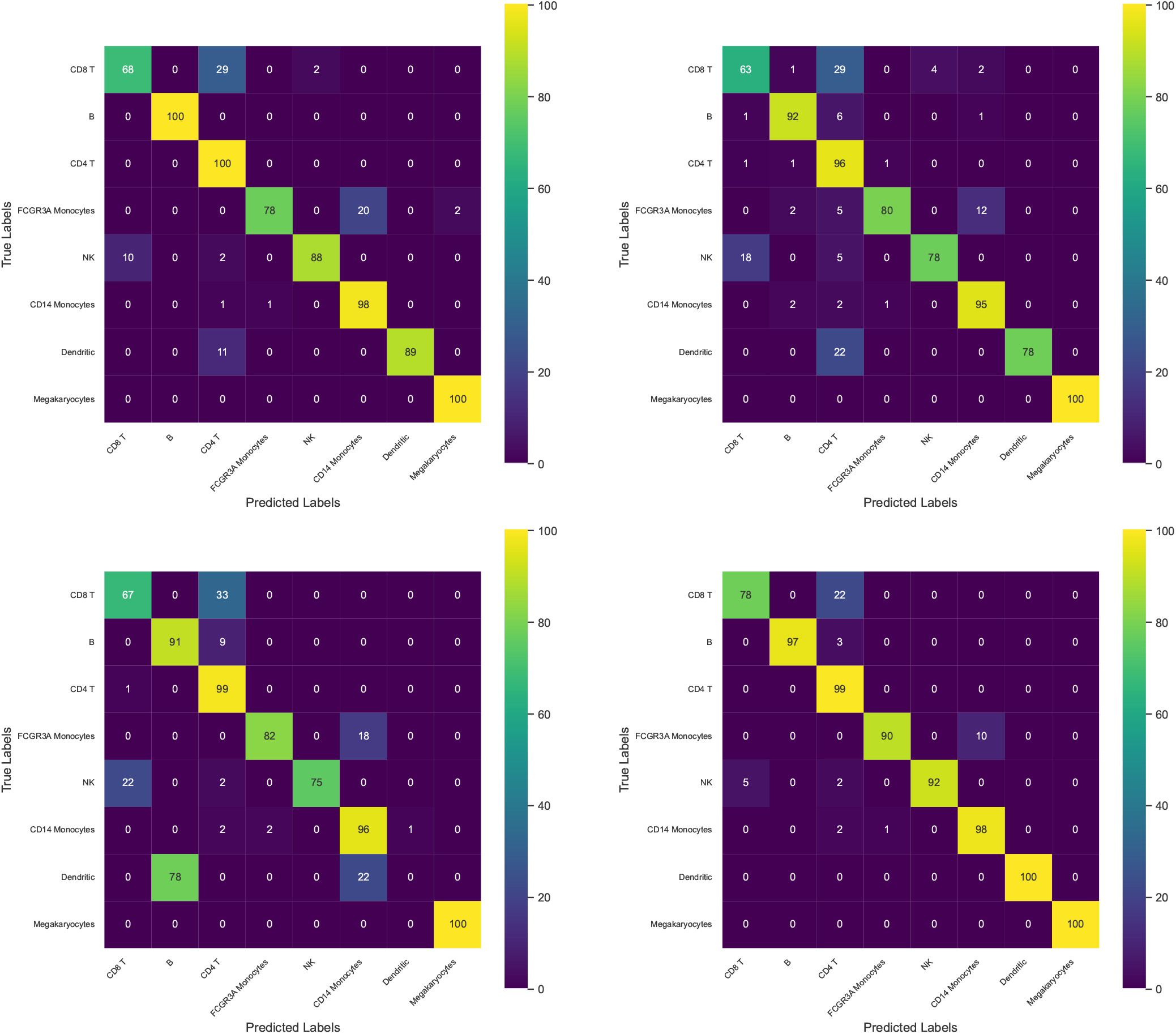
Confusion matrices of (a) scGeneFit, (b) geneBasis, (c) SMaSH and (d) scGIST on the PBMC 3K Dataset for panel size of 60.

**Figure S3:**
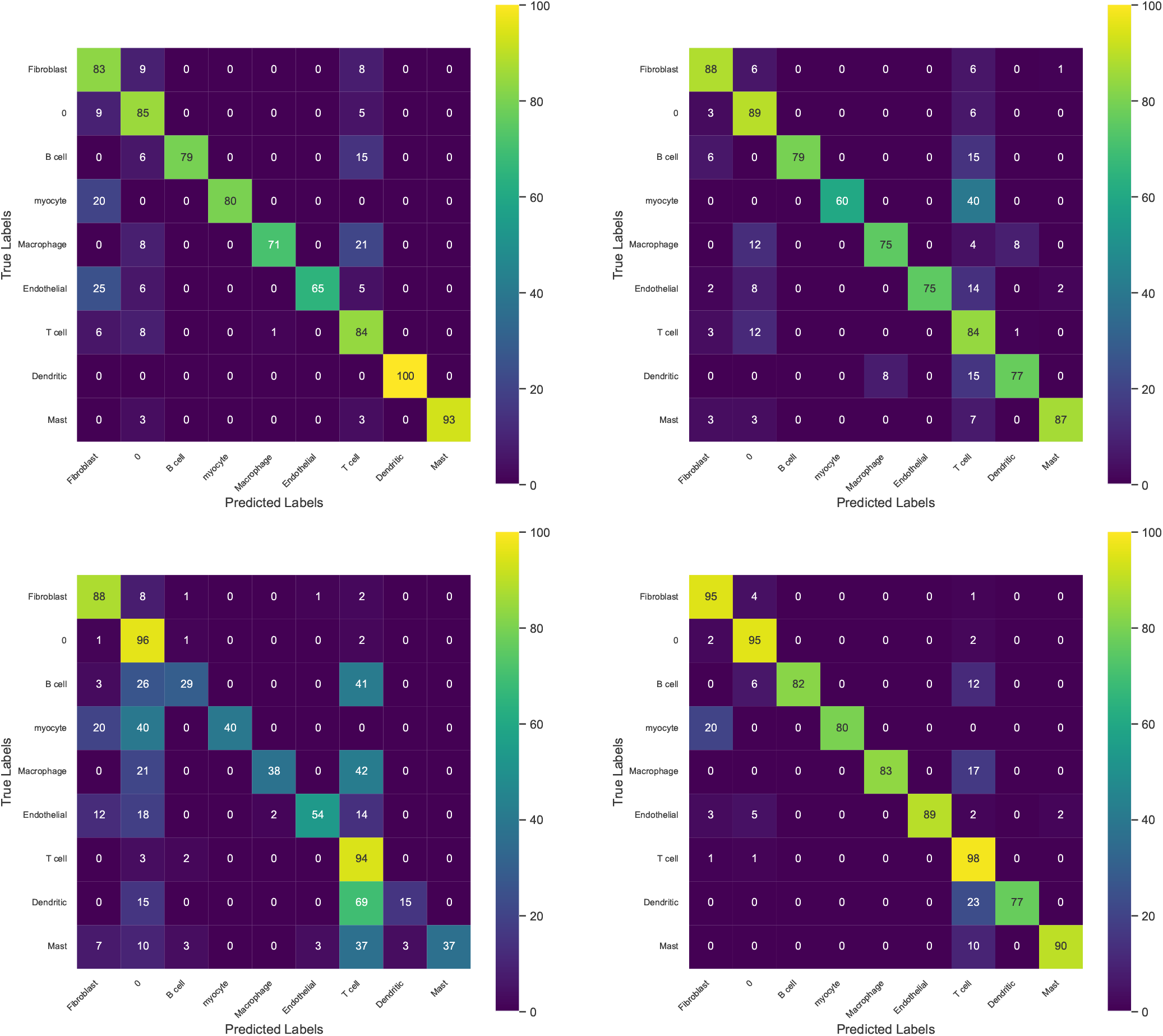
Confusion matrices of (a) scGeneFit, (b) geneBasis, (c) SMaSH and (d) scGIST on the Head and Neck Cancer Dataset for panel size of 60.

**Figure S4:**
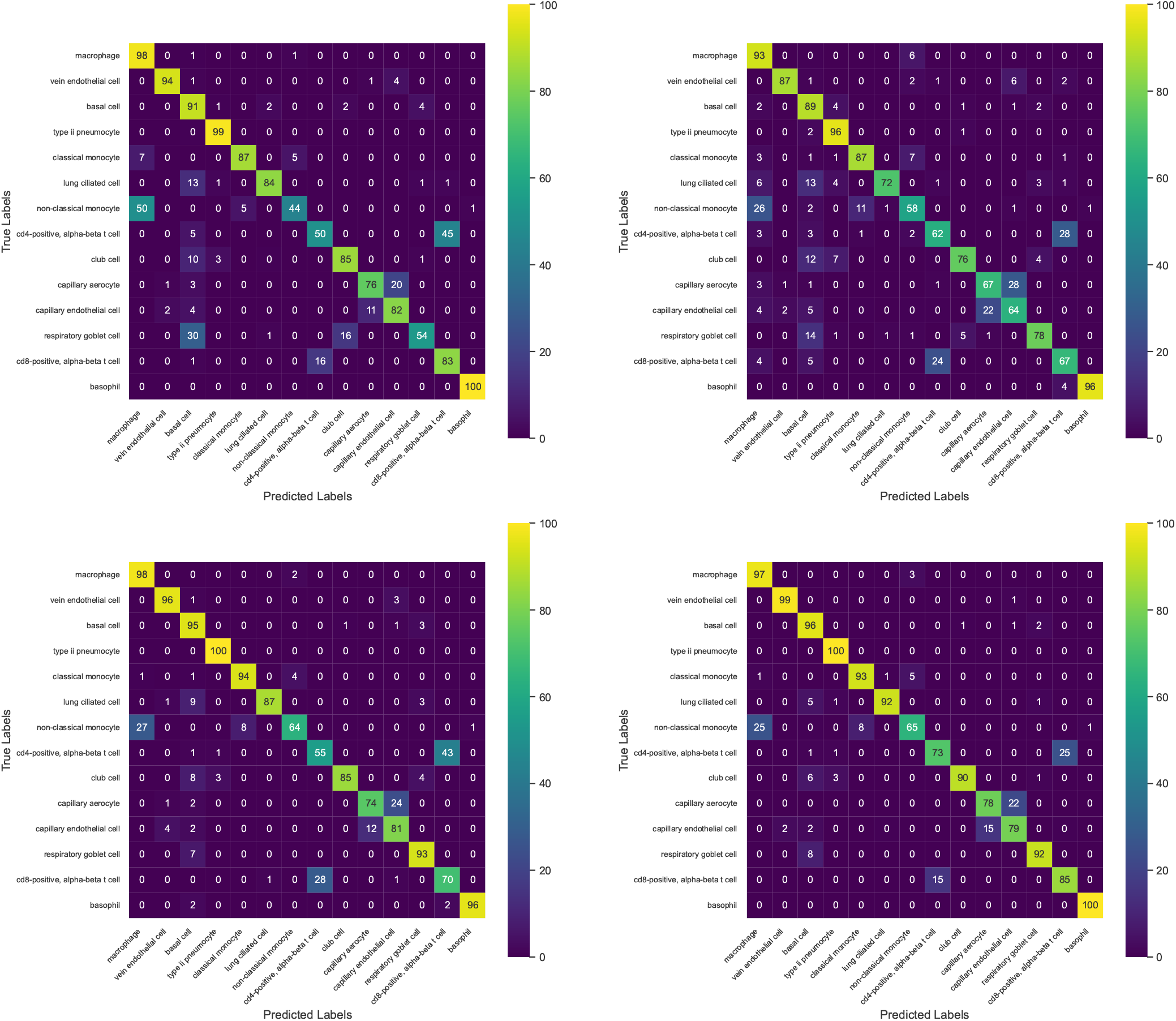
Confusion matrices of (a) scGeneFit, (b) geneBasis, (c) SMaSH and (d) scGIST on the Tabula Sapiens Dataset for panel size of 60.

**Figure S5:**
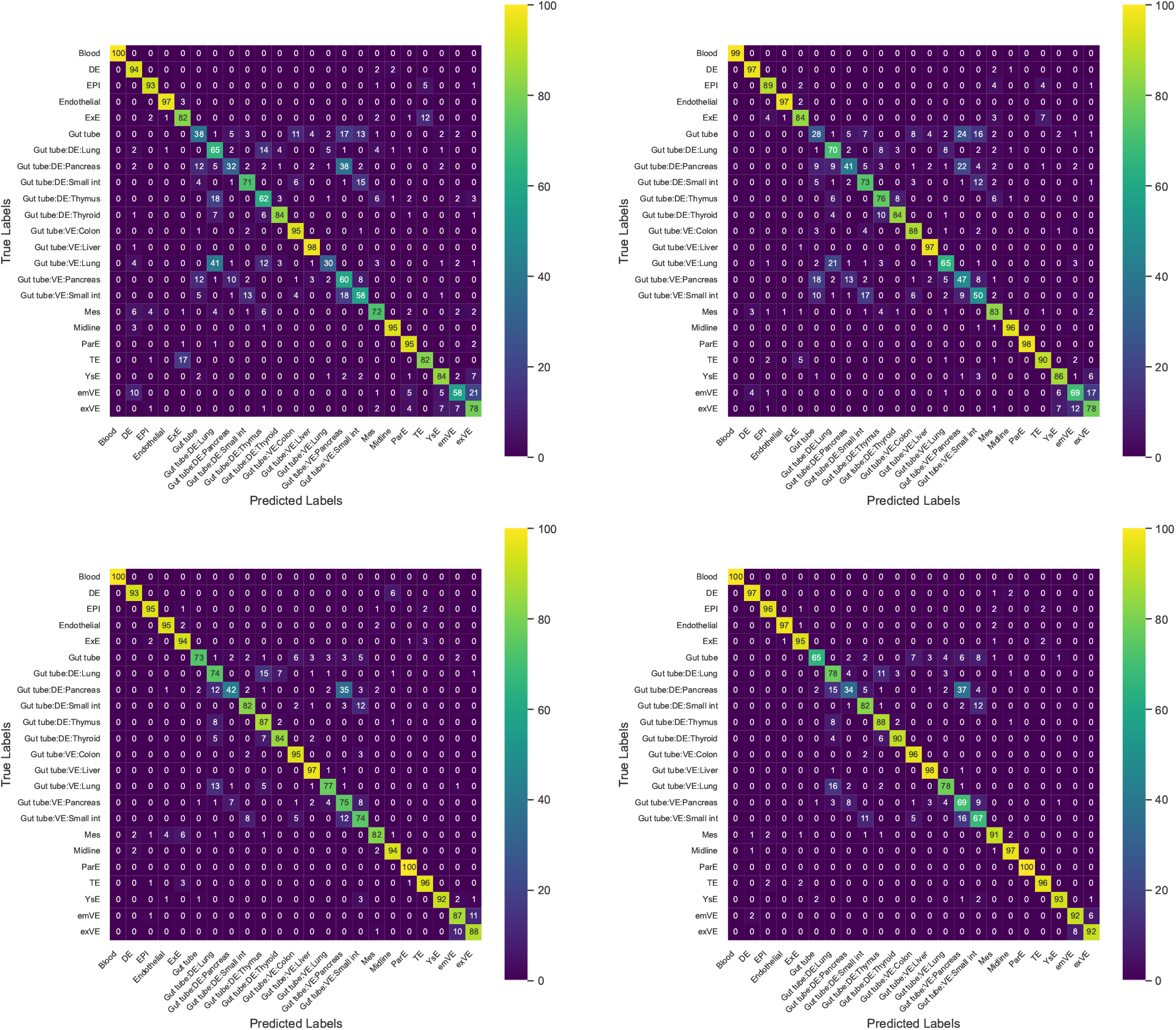
Confusion matrices of (a) scGeneFit, (b) geneBasis, (c) SMaSH and (d) scGIST on the Mouse Endoderm Dataset for panel size of 60.

### 2 Supplementary Tables

**Table S1:**
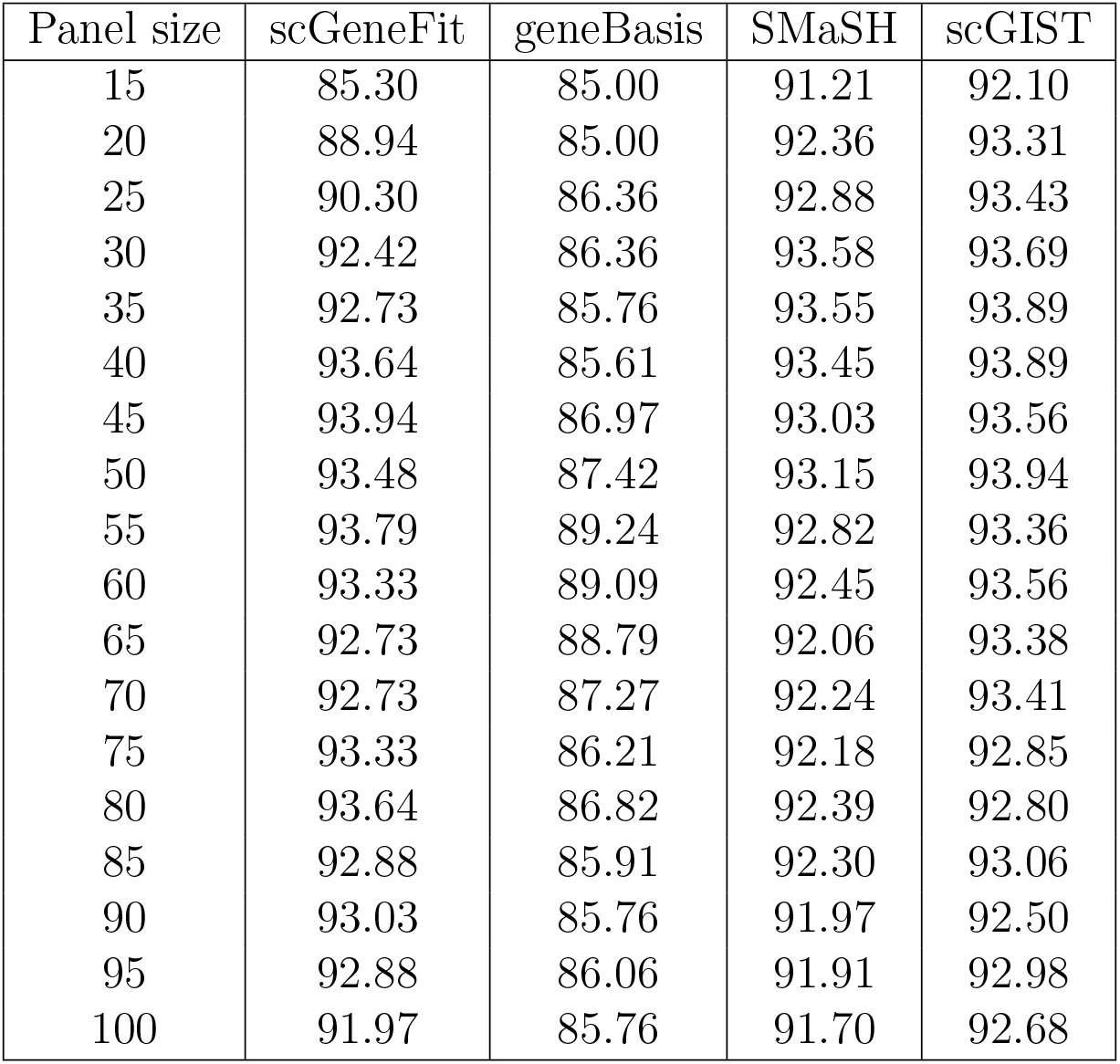
Accuracy of various methods on the PBMC Dataset

| Panel size | scGeneFit | geneBasis | SMaSH | scGIST |
| --- | --- | --- | --- | --- |
| 15 | 85.30 | 85.00 | 91.21 | 92.10 |
| 20 | 88.94 | 85.00 | 92.36 | 93.31 |
| 25 | 90.30 | 86.36 | 92.88 | 93.43 |
| 30 | 92.42 | 86.36 | 93.58 | 93.69 |
| 35 | 92.73 | 85.76 | 93.55 | 93.89 |
| 40 | 93.64 | 85.61 | 93.45 | 93.89 |
| 45 | 93.94 | 86.97 | 93.03 | 93.56 |
| 50 | 93.48 | 87.42 | 93.15 | 93.94 |
| 55 | 93.79 | 89.24 | 92.82 | 93.36 |
| 60 | 93.33 | 89.09 | 92.45 | 93.56 |
| 65 | 92.73 | 88.79 | 92.06 | 93.38 |
| 70 | 92.73 | 87.27 | 92.24 | 93.41 |
| 75 | 93.33 | 86.21 | 92.18 | 92.85 |
| 80 | 93.64 | 86.82 | 92.39 | 92.80 |
| 85 | 92.88 | 85.91 | 92.30 | 93.06 |
| 90 | 93.03 | 85.76 | 91.97 | 92.50 |
| 95 | 92.88 | 86.06 | 91.91 | 92.98 |
| 100 | 91.97 | 85.76 | 91.70 | 92.68 |

**Table S2:** Macro F1-score of various methods on the PBMC Dataset

| Panel size | scGeneFit | geneBasis | SMaSH | scGIST |
| --- | --- | --- | --- | --- |
| 15 | 0.81 | 0.79 | 0.85 | 0.91 |
| 20 | 0.84 | 0.79 | 0.86 | 0.91 |
| 25 | 0.85 | 0.82 | 0.87 | 0.91 |
| 30 | 0.91 | 0.85 | 0.90 | 0.92 |
| 35 | 0.91 | 0.84 | 0.90 | 0.93 |
| 40 | 0.91 | 0.83 | 0.90 | 0.93 |
| 45 | 0.92 | 0.86 | 0.90 | 0.92 |
| 50 | 0.91 | 0.86 | 0.90 | 0.92 |
| 55 | 0.92 | 0.88 | 0.90 | 0.91 |
| 60 | 0.91 | 0.88 | 0.88 | 0.92 |
| 65 | 0.89 | 0.86 | 0.88 | 0.91 |
| 70 | 0.92 | 0.84 | 0.89 | 0.92 |
| 75 | 0.91 | 0.82 | 0.90 | 0.92 |
| 80 | 0.93 | 0.83 | 0.90 | 0.90 |
| 85 | 0.92 | 0.82 | 0.90 | 0.91 |
| 90 | 0.93 | 0.82 | 0.89 | 0.91 |
| 95 | 0.93 | 0.83 | 0.89 | 0.92 |
| 100 | 0.92 | 0.82 | 0.89 | 0.91 |

**Table S3:** Accuracy of various methods on the Head and Neck Cancer Dataset

| Panel size | scGeneFit | geneBasis | SMaSH | scGIST |
| --- | --- | --- | --- | --- |
| 15 | 44.22 | 63.48 | 79.38 | 89.54 |
| 20 | 53.00 | 67.04 | 80.89 | 91.57 |
| 25 | 54.64 | 69.42 | 82.14 | 92.67 |
| 30 | 55.66 | 74.35 | 82.89 | 93.41 |
| 35 | 54.87 | 76.84 | 83.58 | 94.41 |
| 40 | 61.83 | 72.54 | 83.59 | 94.80 |
| 45 | 61.78 | 80.01 | 84.07 | 94.76 |
| 50 | 77.18 | 82.67 | 84.25 | 95.31 |
| 55 | 77.07 | 84.77 | 84.12 | 95.20 |
| 60 | 83.52 | 86.07 | 84.14 | 95.56 |
| 65 | 84.03 | 86.47 | 84.19 | 95.40 |
| 70 | 85.90 | 86.41 | 84.16 | 95.31 |
| 75 | 86.52 | 87.54 | 84.37 | 95.37 |
| 80 | 87.43 | 88.56 | 84.94 | 95.69 |
| 85 | 88.45 | 88.62 | 85.41 | 95.56 |
| 90 | 91.17 | 88.67 | 85.19 | 95.44 |
| 95 | 91.85 | 89.30 | 85.16 | 95.48 |
| 100 | 92.47 | 89.86 | 85.06 | 95.41 |

**Table S4:** Macro F1-score of various methods on the Head and Neck Cancer Dataset

| Panel size | scGeneFit | geneBasis | SMaSH | scGIST |
| --- | --- | --- | --- | --- |
| 15 | 0.69 | 0.53 | 0.40 | 0.81 |
| 20 | 0.70 | 0.61 | 0.44 | 0.87 |
| 25 | 0.74 | 0.64 | 0.46 | 0.89 |
| 30 | 0.75 | 0.75 | 0.50 | 0.89 |
| 35 | 0.74 | 0.76 | 0.51 | 0.91 |
| 40 | 0.74 | 0.75 | 0.52 | 0.92 |
| 45 | 0.74 | 0.78 | 0.53 | 0.92 |
| 50 | 0.80 | 0.79 | 0.54 | 0.92 |
| 55 | 0.79 | 0.81 | 0.53 | 0.92 |
| 60 | 0.83 | 0.81 | 0.55 | 0.93 |
| 65 | 0.83 | 0.83 | 0.54 | 0.92 |
| 70 | 0.75 | 0.84 | 0.56 | 0.92 |
| 75 | 0.74 | 0.85 | 0.57 | 0.92 |
| 80 | 0.78 | 0.86 | 0.58 | 0.93 |
| 85 | 0.78 | 0.86 | 0.58 | 0.93 |
| 90 | 0.80 | 0.87 | 0.57 | 0.93 |
| 95 | 0.80 | 0.88 | 0.57 | 0.93 |
| 100 | 0.80 | 0.88 | 0.58 | 0.92 |

**Table S5:** Accuracy of various methods on the Tabula Sapiens Dataset

| Panel size | scGeneFit | geneBasis | SMaSH | scGIST |
| --- | --- | --- | --- | --- |
| 15 | 43.44 | 72.42 | 84.18 | 86.88 |
| 20 | 51.78 | 77.01 | 86.98 | 89.93 |
| 25 | 62.87 | 78.01 | 87.86 | 91.17 |
| 30 | 71.56 | 79.04 | 88.38 | 91.43 |
| 35 | 84.91 | 82.09 | 88.70 | 91.66 |
| 40 | 87.07 | 82.09 | 89.42 | 91.76 |
| 45 | 86.67 | 82.55 | 90.24 | 91.80 |
| 50 | 87.29 | 83.44 | 90.93 | 92.00 |
| 55 | 88.25 | 84.12 | 91.25 | 92.58 |
| 60 | 88.37 | 85.37 | 91.47 | 92.47 |
| 65 | 88.49 | 85.72 | 91.69 | 92.62 |
| 70 | 87.90 | 86.80 | 91.94 | 92.44 |
| 75 | 88.42 | 86.99 | 91.98 | 92.58 |
| 80 | 88.56 | 87.24 | 92.05 | 92.80 |
| 85 | 88.56 | 86.53 | 92.00 | 92.39 |
| 90 | 88.00 | 86.48 | 91.99 | 92.71 |
| 95 | 88.79 | 86.80 | 91.60 | 92.69 |
| 100 | 88.05 | 86.40 | 91.77 | 92.50 |

**Table S6:**
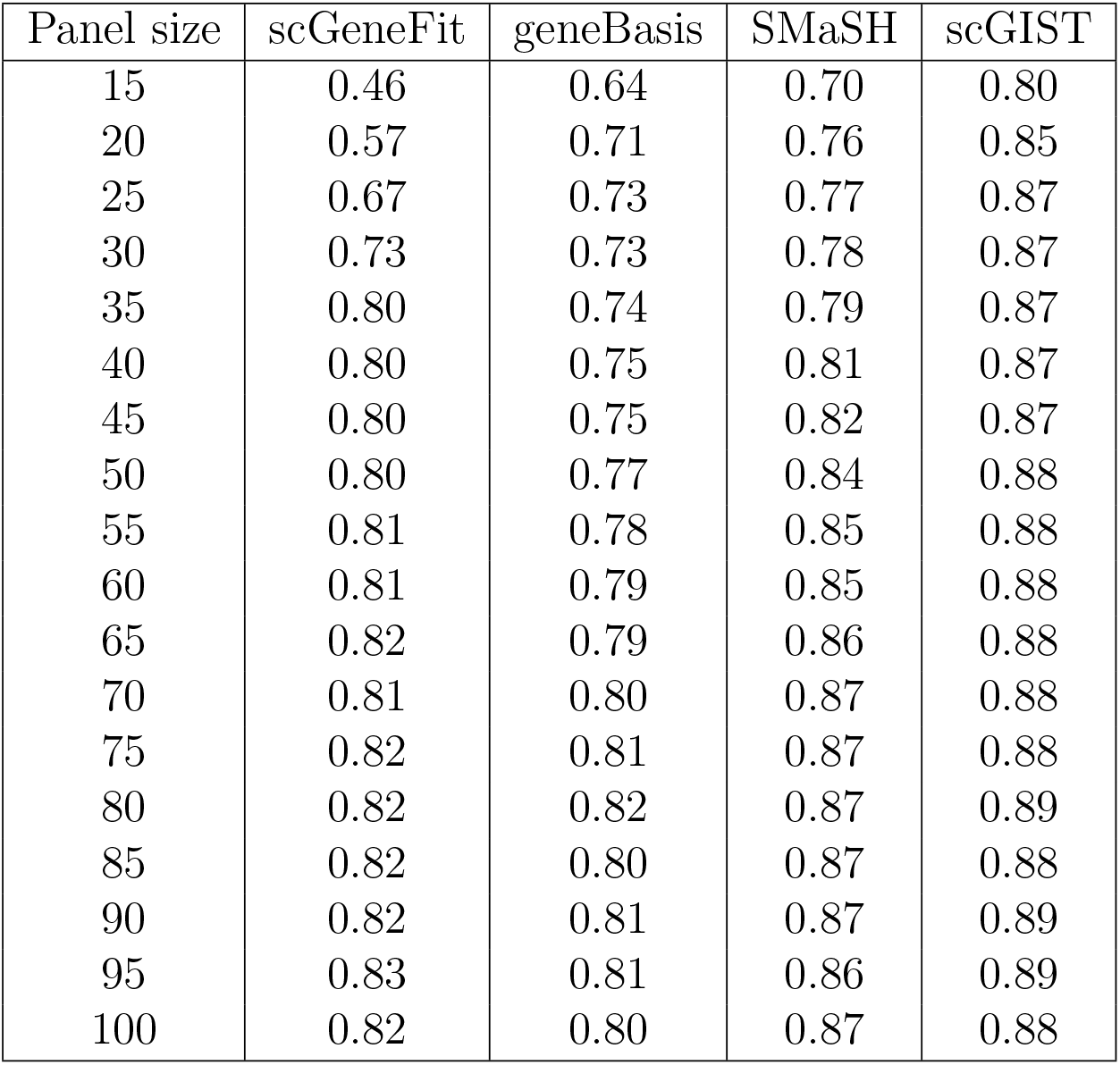
Macro F1-score of various methods on the Tabula Sapiens Dataset

| Panel size | scGeneFit | geneBasis | SMaSH | scGIST |
| --- | --- | --- | --- | --- |
| 15 | 0.46 | 0.64 | 0.70 | 0.80 |
| 20 | 0.57 | 0.71 | 0.76 | 0.85 |
| 25 | 0.67 | 0.73 | 0.77 | 0.87 |
| 30 | 0.73 | 0.73 | 0.78 | 0.87 |
| 35 | 0.80 | 0.74 | 0.79 | 0.87 |
| 40 | 0.80 | 0.75 | 0.81 | 0.87 |
| 45 | 0.80 | 0.75 | 0.82 | 0.87 |
| 50 | 0.80 | 0.77 | 0.84 | 0.88 |
| 55 | 0.81 | 0.78 | 0.85 | 0.88 |
| 60 | 0.81 | 0.79 | 0.85 | 0.88 |
| 65 | 0.82 | 0.79 | 0.86 | 0.88 |
| 70 | 0.81 | 0.80 | 0.87 | 0.88 |
| 75 | 0.82 | 0.81 | 0.87 | 0.88 |
| 80 | 0.82 | 0.82 | 0.87 | 0.89 |
| 85 | 0.82 | 0.80 | 0.87 | 0.88 |
| 90 | 0.82 | 0.81 | 0.87 | 0.89 |
| 95 | 0.83 | 0.81 | 0.86 | 0.89 |
| 100 | 0.82 | 0.80 | 0.87 | 0.88 |

**Table S7:** Accuracy of various methods on the Mouse Endoderm Dataset

| Panel size | scGeneFit | geneBasis | SMaSH | scGIST |
| --- | --- | --- | --- | --- |
| 15 | 42.62 | 56.02 | 67.25 | 77.43 |
| 20 | 43.78 | 61.61 | 73.99 | 80.91 |
| 25 | 45.39 | 66.76 | 78.91 | 84.31 |
| 30 | 52.54 | 69.93 | 81.68 | 85.52 |
| 35 | 58.39 | 71.91 | 83.94 | 86.38 |
| 40 | 66.35 | 72.72 | 85.26 | 87.25 |
| 45 | 68.28 | 74.57 | 86.29 | 87.35 |
| 50 | 73.57 | 76.73 | 86.76 | 88.19 |
| 55 | 75.08 | 77.34 | 87.10 | 88.17 |
| 60 | 75.31 | 77.99 | 87.56 | 88.08 |
| 65 | 76.95 | 79.10 | 87.61 | 88.19 |
| 70 | 78.42 | 79.53 | 88.05 | 88.60 |
| 75 | 79.82 | 79.58 | 88.32 | 88.75 |
| 80 | 80.98 | 80.29 | 88.36 | 88.76 |
| 85 | 80.50 | 80.55 | 88.58 | 88.90 |
| 90 | 81.05 | 81.26 | 88.70 | 88.87 |
| 95 | 81.44 | 81.32 | 88.75 | 89.03 |
| 100 | 81.63 | 81.17 | 89.07 | 88.83 |

**Table S8:** Macro F1-score of various methods on the Mouse Endoderm Dataset

| Panel size | scGeneFit | geneBasis | SMaSH | scGIST |
| --- | --- | --- | --- | --- |
| 15 | 0.41 | 0.55 | 0.66 | 0.77 |
| 20 | 0.42 | 0.60 | 0.73 | 0.80 |
| 25 | 0.44 | 0.66 | 0.78 | 0.84 |
| 30 | 0.51 | 0.69 | 0.81 | 0.85 |
| 35 | 0.57 | 0.71 | 0.83 | 0.86 |
| 40 | 0.65 | 0.72 | 0.85 | 0.87 |
| 45 | 0.67 | 0.74 | 0.86 | 0.87 |
| 50 | 0.72 | 0.76 | 0.86 | 0.87 |
| 55 | 0.74 | 0.76 | 0.86 | 0.87 |
| 60 | 0.74 | 0.77 | 0.87 | 0.87 |
| 65 | 0.76 | 0.78 | 0.87 | 0.87 |
| 70 | 0.77 | 0.79 | 0.87 | 0.88 |
| 75 | 0.79 | 0.78 | 0.88 | 0.88 |
| 80 | 0.80 | 0.79 | 0.88 | 0.88 |
| 85 | 0.79 | 0.80 | 0.88 | 0.88 |
| 90 | 0.80 | 0.80 | 0.88 | 0.88 |
| 95 | 0.80 | 0.80 | 0.88 | 0.88 |
| 100 | 0.80 | 0.80 | 0.88 | 0.88 |

**Table S9:** Accuracy and numbers of genes with easy to design probes included in the panel by scGIST when no genes are assigned priority and *β* = 0.2

| Panel size | Accuracy | No. genes with easy to design probes taken |
| --- | --- | --- |
| 50 | 87.52 | 17 |
| 75 | 88.87 | 32 |
| 100 | 89.26 | 36 |
| 125 | 89.40 | 46 |
| 150 | 89.76 | 46 |
| 175 | 88.75 | 57 |
| 200 | 89.41 | 64 |
| 225 | 89.70 | 73 |
| 250 | 88.55 | 83 |
| 275 | 89.11 | 84 |
| 300 | 89.14 | 86 |
| 325 | 88.91 | 95 |
| 350 | 88.39 | 108 |
| 375 | 88.78 | 110 |
| 400 | 88.94 | 117 |
| 425 | 88.87 | 119 |
| 450 | 89.05 | 116 |
| 475 | 88.67 | 124 |
| 500 | 88.81 | 133 |

**Table S10:** Accuracy and numbers of genes with easy to design probes included in the panel by scGIST when genes with easy to design probes are assigned priority score 1 and *β* = 0.2

| Panel size | Accuracy | No. genes with easy to design probes taken |
| --- | --- | --- |
| 50 | 87.16 | 31 |
| 75 | 88.69 | 43 |
| 100 | 88.64 | 60 |
| 125 | 88.22 | 76 |
| 150 | 88.34 | 93 |
| 175 | 88.66 | 110 |
| 200 | 88.57 | 131 |
| 225 | 89.28 | 151 |
| 250 | 88.72 | 172 |
| 275 | 88.43 | 195 |
| 300 | 88.40 | 212 |
| 325 | 88.19 | 240 |
| 350 | 87.69 | 258 |
| 375 | 87.31 | 286 |
| 400 | 86.80 | 312 |
| 425 | 86.15 | 345 |
| 450 | 85.85 | 366 |
| 475 | 85.71 | 385 |
| 500 | 85.92 | 385 |

**Table S11:** Accuracy and numbers of genes with easy to design probes included in the panel by scGIST for varying *β* when genes with easy to design probes are assigned priority score 1 and panel size is 400

| Beta ($\beta$) | Accuracy | No. genes with easy to design probes taken |
| --- | --- | --- |
| 0 | 88.93 | 114 |
| 0.025 | 88.78 | 135 |
| 0.05 | 89.17 | 161 |
| 0.075 | 89.31 | 187 |
| 0.1 | 88.84 | 220 |
| 0.125 | 87.87 | 244 |
| 0.15 | 87.21 | 266 |
| 0.175 | 87.43 | 289 |
| 0.2 | 86.57 | 315 |
| 0.225 | 85.74 | 330 |
| 0.25 | 85.39 | 348 |
| 0.275 | 84.81 | 357 |
| 0.3 | 84.96 | 362 |
| 0.325 | 83.73 | 373 |
| 0.35 | 83.61 | 376 |
| 0.375 | 83.43 | 382 |
| 0.4 | 82.84 | 385 |
| 0.425 | 82.59 | 385 |
| 0.45 | 82.72 | 385 |
| 0.475 | 83.04 | 385 |
| 0.5 | 82.65 | 385 |

### 3 Supplementary Notes

#### 3.1 Alternate regularization penalty for panel size

In the default configuration, scGIST is strict about panel size, which means the regularization function penalizes for including more or less genes than the given panel size. It may not always be the desired outcome. Sometimes we want to create a panel with as few genes as possible without sacrificing accuracy. In the alternate regularization function, we change the second part of the regularization function to *α* max(||*w*||_1_ − *d*, 0) from *α*|||*w*||_1_−*d*|. Supplementary Figure S6 shows the regularization loss against number of genes included in the panel for default and alternate regularization functions. The accracy and the numbers of genes included in the panel for varying target panel size for the two modes are shown in Supplementary Figure S7.

**Figure S6:**
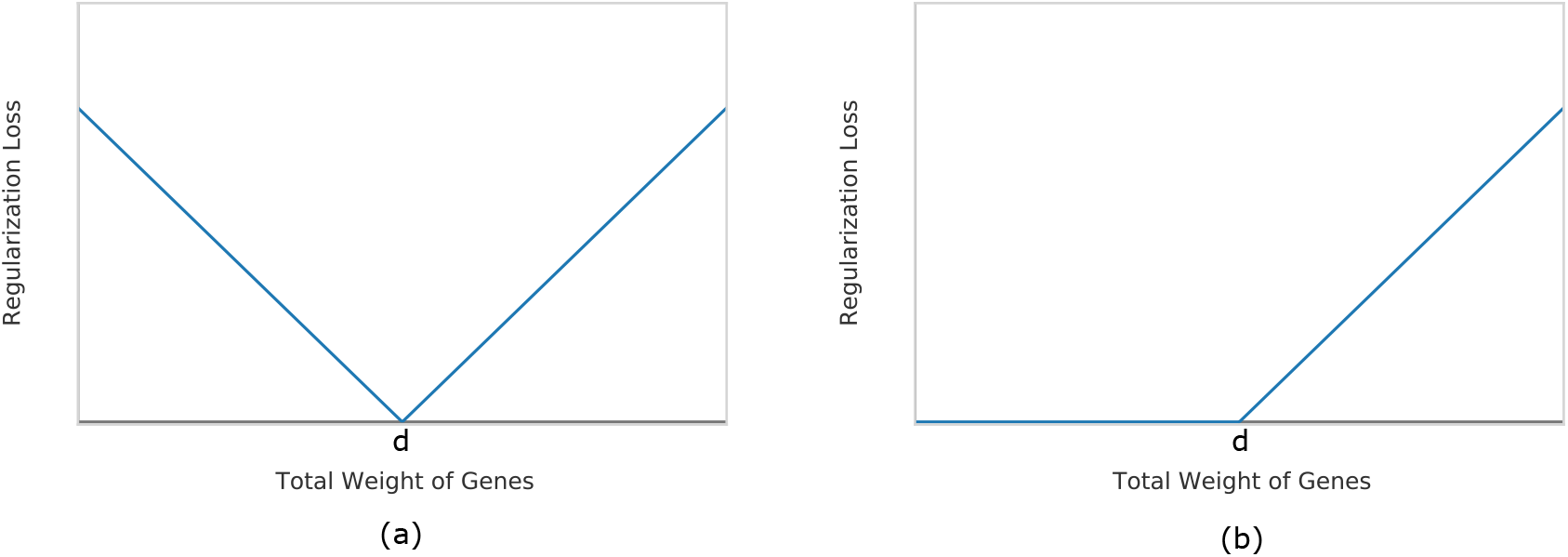
Comparison of regularization loss vs number of genes included in panels for (a) default and (b) alternate regularization functions.

**Figure S7:**
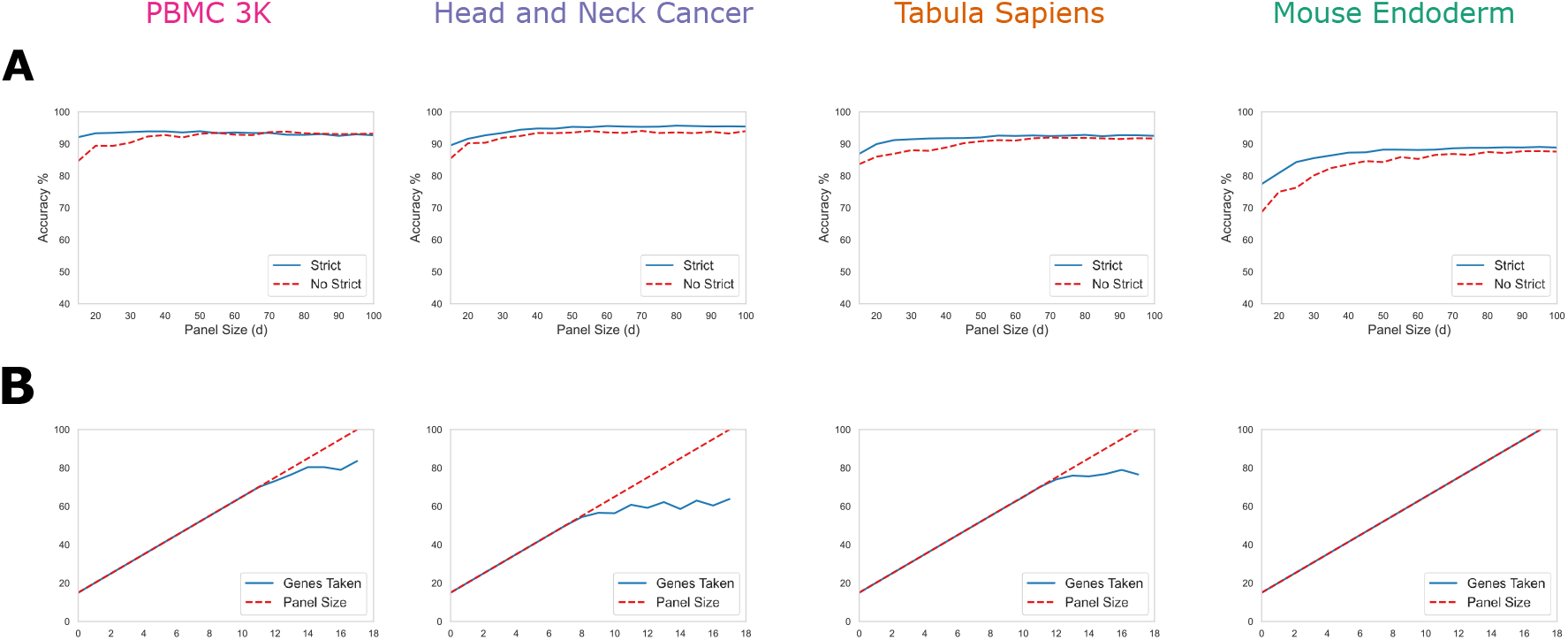
**A.** Accuracy of the default (strict) mode and the alternate (non-strict) mode for different datasets for varying target panel size, **B.** Comparison of target panel size and the numbers of genes included in the panel in for the two modes.

#### 3.2 Evaluation metrics

Cell type prediction is multi-class classification problem with possible class imbalance. We used accuracy and macro F1-score as the evaluation metrics to assess various tools. Consider, a dataset with *N* samples and *K* possibles classes. Let, *TP*_*k*_, *TN*_*k*_, *TP*_*k*_, *TN*_*k*_ be the number of true positives, true negatives, false positives and false negatives for the *k*-the class type. Then the accuracy is given by

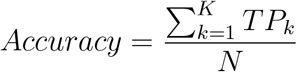

However, accuracy can be misleading when there is class imbalance. So, we also use macro F1-score for assessment.

Precision and Recall for the *k*-th class type are given by

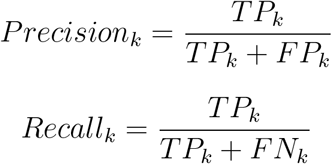

The macro average precision and recall are the averages over the precisions and recalls.

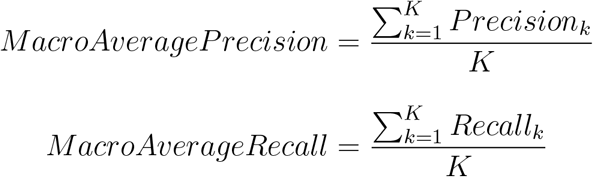

The macro F1-score is then given by

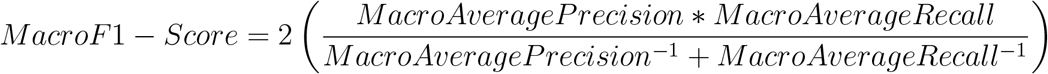

**References**

## References

1. Longo, S. K., Guo, M. G., Ji, A. L. & Khavari, P. A. Integrating single-cell and spatial transcriptomics to elucidate intercellular tissue dynamics. Nat. Rev. Genet. 22, 627–644 (2021).

2. Rao, A., Barkley, D., França, G. S. & Yanai, I. Exploring tissue architecture using spatial transcriptomics. Nature 596, 211–220 (2021).

3. Moffitt, J. R., Lundberg, E. & Heyn, H. The emerging landscape of spatial profiling technologies. Nat. Rev. Genet. 1–19 (2022).

4. Moses, L. & Pachter, L. Museum of spatial transcriptomics. Nat. Methods 19, 534–546 (2022).

5. Canozo, F. J. G., Zuo, Z., Martin, J. F. & Samee, M. A. H. Cell-type modeling in spatial transcriptomics data elucidates spatially variable colocalization and communication between cell-types in mouse brain. Cell Syst. 13, 58–70 (2022).

6. Cleary, B. et al. Compressed sensing for highly efficient imaging transcriptomics. Nat. Biotechnol. 39, 936–942 (2021).

7. Levsky, J. M., Shenoy, S. M., Pezo, R. C. & Singer, R. H. Single-cell gene expression profiling. Science 297, 836– 840 (2002).

8. Lubeck, E. & Cai, L. Single-cell systems biology by super-resolution imaging and combinatorial labeling. Nat. methods 9, 743–748 (2012).

9. Moffitt, J. R. et al. Molecular, spatial, and functional single-cell profiling of the hypothalamic preoptic region. Science 362, eaau5324 (2018).

10. Zhang, M. et al. Spatially resolved cell atlas of the mouse primary motor cortex by merfish. Nature 598, 137–143 (2021).

11. Hara, T. et al. Interactions between cancer cells and immune cells drive transitions to mesenchymal-like states in glioblastoma. Cancer Cell 39, 779–792 (2021).

12. Lu, Y. et al. Spatial transcriptome profiling by merfish reveals fetal liver hematopoietic stem cell niche architecture. Cell Discov. 7, 1–17 (2021).

13. Li, Y., Chen, C.-Y. & Wasserman, W. W. Deep feature selection: theory and application to identify enhancers and promoters. J. Comput. Biol. 23, 322–336 (2016).

14. Dumitrascu, B., Villar, S., Mixon, D. G. & Engelhardt, B. E. Optimal marker gene selection for cell type discrimination in single cell analyses. Nat. communications 12, 1–8 (2021).

15. Missarova, A. et al. geneBasis: an iterative approach for unsupervised selection of targeted gene panels from scRNA-seq. Genome biology 22, 1–22 (2021).

16. Nelson, M. E., Riva, S. G. & Cvejic, A. SMaSH: a scalable, general marker gene identification framework for single-cell rna-sequencing. BMC Bioinforma. 23, 1– 16 (2022).

17. Stoeckius, M. et al. Simultaneous epitope and transcriptome measurement in single cells. Nat. methods 14, 865– 868 (2017).

18. Puram, S. V. et al. Single-cell transcriptomic analysis of primary and metastatic tumor ecosystems in head and neck cancer. Cell 171, 1611–1624 (2017).

19. Consortium, T. S.* et al. The tabula sapiens: A multiple-organ, single-cell transcriptomic atlas of humans. Science 376, eabl4896 (2022).

20. Nowotschin, S. et al. The emergent landscape of the mouse gut endoderm at single-cell resolution. Nature 569, 361–367 (2019).

21. Van der Maaten, L. & Hinton, G. Visualizing data using t-sne. J. machine learning research 9 (2008).

22. Lohoff, T. et al. Integration of spatial and single-cell transcriptomic data elucidates mouse organogenesis. Nat. biotechnology 40, 74–85 (2022).

23. Pijuan-Sala, B. et al. A single-cell molecular map of mouse gastrulation and early organogenesis. Nature 566, 490–495 (2019).

24. Xiao, Y. et al. Hippo signaling plays an essential role in cell state transitions during cardiac fibroblast development. Dev. cell 45, 153–169 (2018).

25. Calvo, F. et al. Mechanotransduction and yap-dependent matrix remodelling is required for the generation and maintenance of cancer-associated fibroblasts. Nat. cell biology 15, 637–646 (2013).

26. Shin, E. & Kim, J. The potential role of yap in head and neck squamous cell carcinoma. Exp. & Mol. Medicine 52, 1264–1274 (2020).

27. Traag, V. A., Waltman, L. & Van Eck, N. J. From louvain to leiden: guaranteeing well-connected communities. Sci. reports 9, 1–12 (2019).

28. Armingol, E., Officer, A., Harismendy, O. & Lewis, N. E. Deciphering cell–cell interactions and communication from gene expression. Nat. Rev. Genet. 22, 71–88 (2021).

29. Efremova, M., Vento-Tormo, M., Teichmann, S. A. & Vento-Tormo, R. Cellphonedb: inferring cell–cell communication from combined expression of multi-subunit ligand–receptor complexes. Nat. protocols 15, 1484– 1506 (2020).

30. Cleary, B., Cong, L., Cheung, A., Lander, E. S. & Regev, Efficient generation of transcriptomic profiles by random composite measurements. Cell 171, 1424–1436 (2017).

31. Wolf, F. A., Angerer, P. & Theis, F. J. Scanpy: largescale single-cell gene expression data analysis. Genome biology 19, 1–5 (2018).

32. Satija, R., Farrell, J. A., Gennert, D., Schier, A. F. & Regev, A. Spatial reconstruction of single-cell gene expression data. Nat. biotechnology 33, 495–502 (2015).

